# SimdMinimizers: Computing random minimizers, *fast*

**DOI:** 10.1101/2025.01.27.634998

**Authors:** Ragnar Groot Koerkamp, Igor Martayan

## Abstract

**Motivation:** Because of the rapidly-growing amount of sequencing data, computing *sketches* of large textual datasets has become an essential preprocessing task. These sketches are typically much smaller than the input sequences, but preserve sufficient information for downstream analysis. *Minimizers* are an especially popular sketching technique and used in a wide variety of applications. They sample at least one out of every *w* consecutive *k*-mers. As DNA sequencers are getting more accurate, some applications can afford to use a larger *w* and hence sparser and smaller sketches. And as sketches get smaller, their analysis becomes faster, so the time spent sketching the full-sized input becomes more of a bottleneck.

**Methods:** Our library `simd-minimizers` implements a random minimizer algorithm using SIMD instructions. It supports both AVX2 and NEON architectures. Its main novelty is two-fold. First, it splits the input into 8 chunks that are streamed over in parallel through all steps of the algorithm. This is enabled by using the completely deterministic *two-stacks* sliding window minimum algorithm, which seems not to have been used before for finding minimizers.

**Results:** Our library is up to 6.8× faster than a scalar implementation of the *rescan* method when *w* = 5 is small, and 3.4× faster for larger *w* = 19. Computing *canonical* minimizers is less than 50% slower than computing forward minimizers, and over 15× faster than the existing implementation in the `minimizer-iter` crate. Our library finds all (canonical) minimizers of a 3.2 Gbp human genome in 5.2 (resp. 6.7) seconds.

## 1 Introduction

*Minimizers* were simultaneously introduced by [29] and [28] as a method to sample short strings of fixed length *k*, called *k*-mers or *k*-grams, for the purpose of fingerprinting and comparing large textual documents such as genomic sequences. This sampling method plays a central role in bioinformatics for the high-throughput analysis of DNA sequencing data and is a fundamental building block for many related tasks such as indexing [27, 19], counting [3, 21], aligning [17, 12], or assembling [4, 1] genomic sequences.

Minimizers are defined as follows: given a window *W* of *w* consecutive *k*-mers, the minimizer of *W* is the smallest *k*-mer according to some order. In practice, this order is often pseudo-random by hashing the *k*-mers, leading to the *random minimizer*. The *density* of a minimizer scheme is the expected fraction of sampled *k*-mers on a sufficiently long random string. When *k* is not too small, random minimizers have an expected density close to 2*/*(*w* + 1), which is around twice as much as the lower bound of 1*/w*. In recent years, there have been a number of papers on methods with lower density than random minimizers [33, 26, 9, 7, 6]. While these contributions have narrowed the gap to an optimal-density sampling scheme [15], none of them focused on improving the computation time of minimizers.

In bioinformatics applications, the *k*-mer length is usually at most *k* ≤ 31, so that each *k*-mer can be represented by a single `u64` machine word, and the window length can be as low as 5 (where around a third of the *k*-mers is sampled), but is typically between 10 and 30. Longer windows of size up to 100 are also possible to index highly conserved texts.

### Problem statement

We aim to solve the following problem as fast as possible: given a bitpacked representation of a sequence of ACGT DNA characters, compute the positions of all (canonical) random minimizers.

### Contributions

This work introduces a carefully optimized algorithm to compute the minimizers of a genomic sequence. Conceptually, it consists of two parts. First, we introduce a scalar algorithm (Section 3) that uses *O*(*n*) time and *O*(*w*) space. Most of this can be trivially parallelized to *L* = 8 independent lanes with AVX2 or NEON SIMD instructions, and in Section 4 we specifically handle the input and output. The parts we discuss are:

1. We apply ntHash [23, 14], a pseudo-random rolling hash function for *k*-mers (Section 3.2).
2. We compute the sliding-window minima of the hashes using the *two-stacks* method [10, 30] (Section 3.3).
3. We compute canonical minimizers based on *refined minimizers* [25] to decide the *strand* of each window (Section 3.4).
4. We extend the scalar algorithm to SIMD by using it on *L* chunks of the (bitpacked) input sequence in parallel (Section 4.1).
5. Lastly, we collect and deduplicate the *L* parallel streams into unique minimizer positions (Section 4.2).

### Results

Our method is 3.4× (for large *w* = 19) to 6.8× (for small *w* = 5) times faster than the fastest non-SIMD algorithm for computing forward minimizers. For canonical minimizers, we only compare against a simple implementation and find over 15 × speedup. As a result, we can compute the minimizers of a human genome in 5.2 seconds, and the canonical minimizers in 6.7 seconds. We also adapt our method to support generic plain-text ASCII input (|Σ| = 256), which is slightly (30%) slower due to the larger input characters.

### Software

A Rust implementation of our method is publicly available at https://github.com/rust-seq/simd-minimizers. The packed sequence representation, the splitting into chunks, and the parallel iteration over their bases is extracted to a separate library, packed-seq, available at https://github.com/rust-seq/packed-seq.

## 2 Preliminaries

### Bitpacking

In this work, we assume that the input sequence is over the DNA alphabet Σ = {A, C, T, G} and that each letter is encoded using two bits: A = 00, C = 01, T = 10, G = 11. This encoding can easily be obtained from the ASCII representation by applying a mask: (c >> 1) & 3. Additionally, we assume that the whole sequence is bitpacked using this 2-bit encoding, which can be done as a preprocessing step on the input if necessary. Non-ACGT characters have to be handled during this preprocessing as well, and could be skipped, converted to A, or the input sequence could be split at these points. We assume that the hardware is little-endian, and that the integer value of a sequence *x*_0_*x*_1_*x*_2_ … *x*_*k−*1_ is given by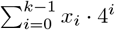

### Minimizers

Given parameters *w* and *k*, a *window W* of length *𝓁* = *w* + *k* − 1 contains *w* consecutive *k*-mers. The *minimizer* of the window is the smallest *k*-mer in the window. For *random* minimizers, *k*-mers are ordered by a pseudo-random order, usually given by comparing hashes of the *k*-mers. In case of ties, the leftmost smallest *k*-mer is chosen.

Our goal is to compute the absolute position of the minimizer of every window *W* in the input text. Since adjacent windows often have the same *k*-mer as minimizer, we only want each position to be listed once in the output.

### Canonical minimizers

Because DNA is double-stranded and most sequencing technologies do not distinguish these two strands, genomic sequences have an additional constraint: a sequence and its *reverse-complement* (the reversed sequence of complementary bases `A ↔ T` and `C ↔ G`) should be considered identical. To satisfy this constraint, *canonical minimizers* should return the same set of *k*-mers regardless of the strandedness of the input. Specifically, if the canonical minimizer of a window *W* is at position *p*, then the canonical minimizer of the reverse-complement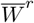of *W* should be at position |*W*| − *k* − *p* = *w* − 1 − *p*. In practice, canonical minimizers are often overlooked as an implementation detail and most existing methods simply compare canonical *k*-mers, computed as *x*^*c*^ = min(*x, x*^*r*^), which gives a weaker guarantee [22].

## 3 A predictable scalar algorithm

### Overview

Computing the minimizers of a sequence generally involves a few steps, as shown in Figure 1. First, the *k*-mers are hashed. We do this using ntHash, a rolling hash (Section 3.2). Next, in each window of *w k*-mer hashes, we must find the position of the leftmost *k*-mer with the smallest hash. For this, we use a method based on the two-stacks algorithm (Section 3.3). Lastly, many adjacent windows will have the same minimizer, and thus these positions must be deduplicated.

**Figure 1.**
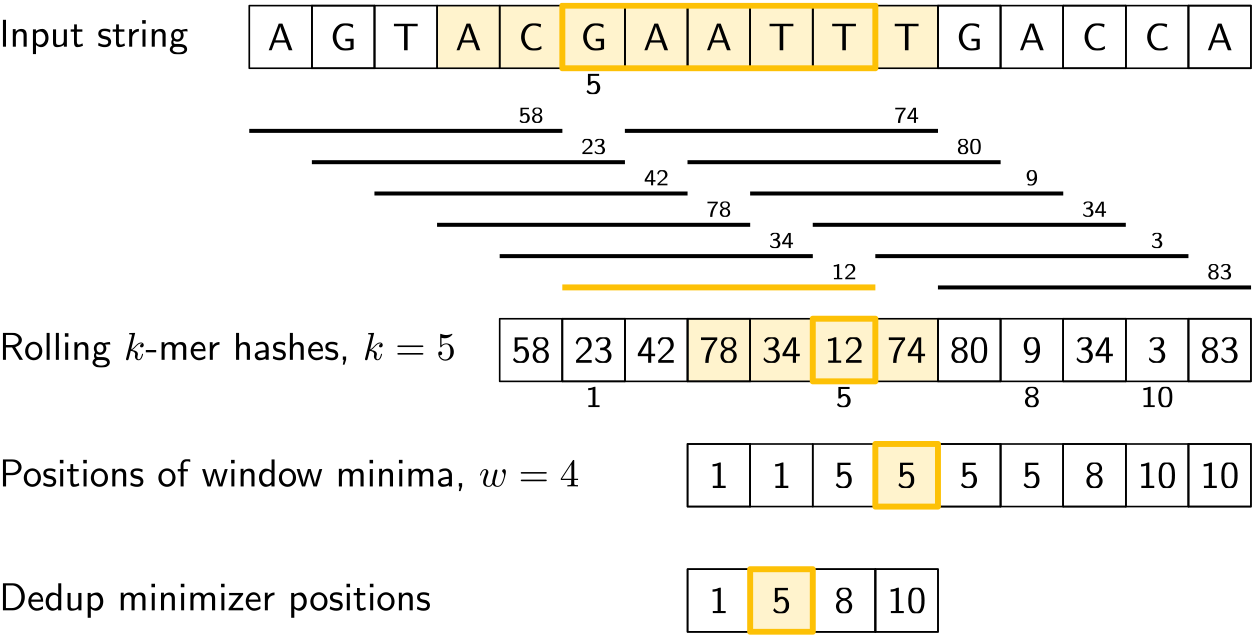
Overview of computing minimizers, for *k* = 5 and *w* = 4. First, all *k*-mers of the input string (shown as black horizontal lines) are hashed (the small numbers above them). Then, for each window of *w* = 4 consecutive *k*-mers, we find the absolute position of its smallest *k*-mer. For example, for the first window, the *k*-mer at position 1 (zero-based) has the smallest hash (23). Similarly, the fourth window (highlighted, spanning *𝓁* = *w* + *k* − 1 = 8 bases) has minimal hash 12 by the *k*-mer at position 5. Lastly, these minimizer positions are deduplicated.

In this section, we present a scalar algorithm (that will be trivial to parallelize later by using SIMD instructions Section 4), and we also introduce a variant to compute canonical minimizers in Section 3.4.

The rolling hash takes constant time per character, and our sliding window algorithm will also take amortized constant time per character, so that the entire method runs in *O*(*n*). It needs *O*(*w*) space to store the hashes of the current window. With the SIMD optimizations, this improves to *O*(*n/L*) time and *O*(*Lw*) space, where *L* is the number of SIMD lanes.

### 3.1 Iterating Packed Input

As input to our algorithm, we use a 2-bit packed sequence representation. Our representation is little-endian, in that the two least significant bits of each 8-bit or 32-bit value correspond to the leftmost encoded base.

#### Extracting bases

To iterate the bases of the input string, we process them one u32 of 16 bases at a time. For each of those, we do 16 iterations where we extract the 2 least significant bits to return, and then shift the remainder down by 2 bits.

For 8-bit ASCII input, instead, the low 8 bits can be extracted and then shifted away.

#### Delayed iterator

In order to support a rolling hash, we simply iterate the characters twice in parallel, with an offset: once for the character *entering* each *k*-mer and once delayed by *k* −1 iterations for the character *leaving* each *k*-mer. Similarly, for canonical minimizers we will also need the character leaving the *window*, for which we do a third iteration delayed by *𝓁* − 1 steps. To avoid additional reads from memory, we store `u32` values read from memory in a small ring buffer.

### 3.2 Rolling Hash

The first step of the algorithm is the computation of a hash for each *k*-mer. We adapt ntHash [23, 14], a popular rolling hash function for DNA sequences that is based on cyclic polynomials [2] as an improvement over Karp-Rabin hashing [13]. This rolling hash function can be summarized as follows: each of the 4 bases is associated to a fixed 32-bit random value, denoted *f* (*x*), and the hash of a *k*-mer *u* = *x*_0_ … *x*_*k−*1_ is computed as 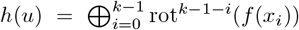, where rot^*i*^ denotes a cyclic rotation by *i* bits to the left, and ⊕ denotes xor. In practice, each *x*_*i*_ is a value in {0, 1, 2, 3} and *f* (*x*_*i*_) is a simple table lookup as shown in Algorithm 2.

#### Algorithm 1

Pseudocode for iterating the packed input sequence, and for iterating over the last (*in*) and first (*out*) bases of all *k*-mers. The code samples are written in terms of “iterator adapters”, where the input is a stream over something, and the output is a transformed stream of *yielded* elements.

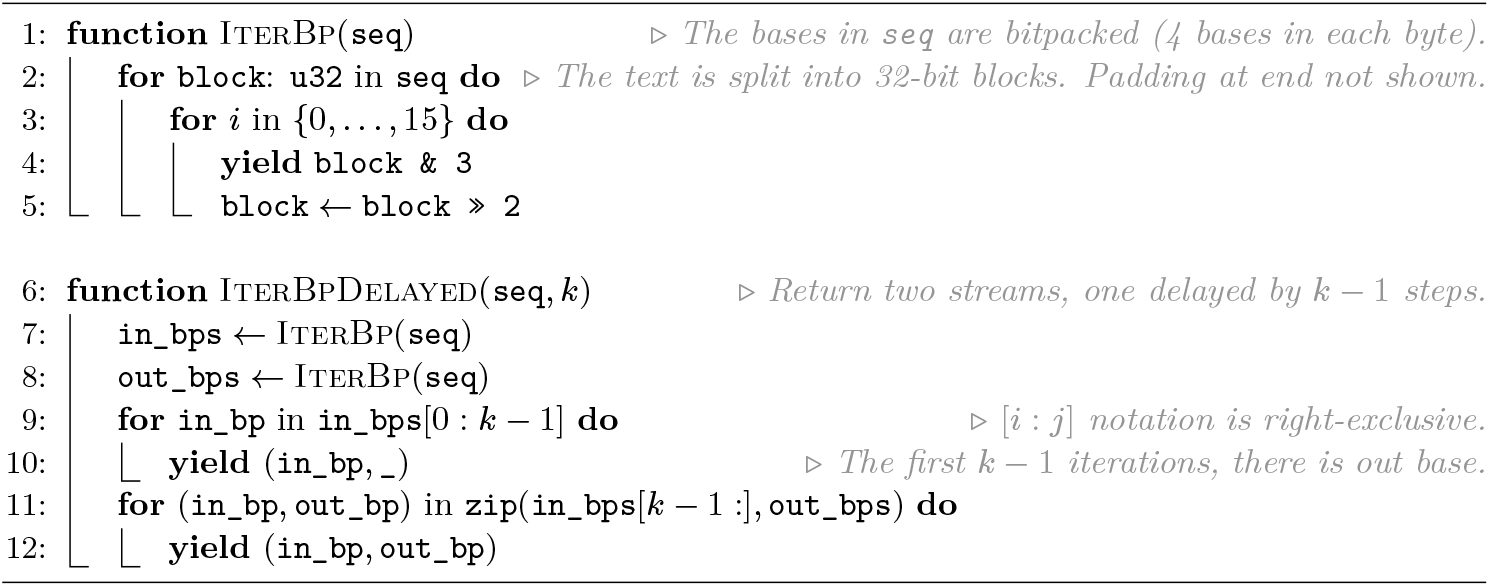

#### Algorithm 2

NtHash rolling hash. The input is an iterator over pairs of characters coming in and out of each *k*-mer as returned by IterBpDelayed (Algorithm 1). The output is an iterator over the hashes of all *k*-mers. The algorithm first processes the first *k* − 1 characters to initialize the rolling hash, and then repeatedly adds and removes one character at a time. Each operation can be vectorized to work on *L* lanes in parallel.

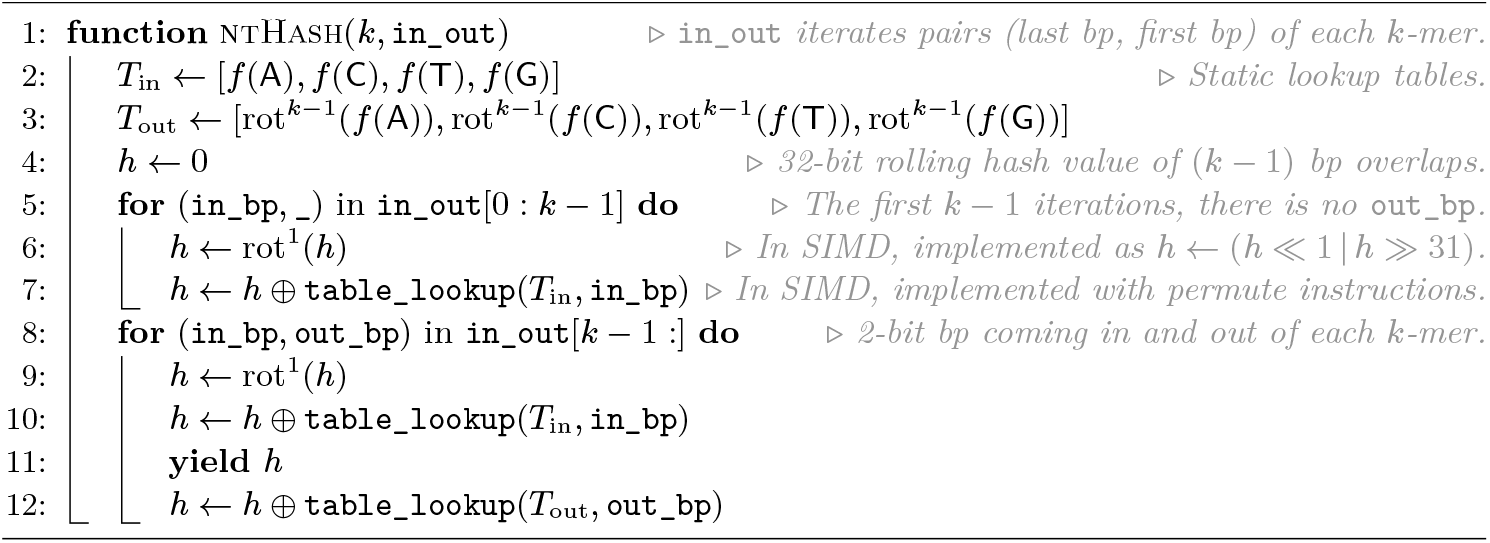

Using a rolling hash has a twofold advantage in our use case: first, we only need the first and last bases of each *k*-mer to update its hash based on the previous one, so that we do not have to store the *k*-mer itself, and second, we are not limited to *k*-mers that fit in a constant number of words.

Most of the pseudocode in Algorithm 2 can be trivially adapted to work on u32×8 SIMD registers containing 8 32-bit values. The `table_lookup` function computes the values of *f* (*x*) of the *L* nucleotides stored in `in_bp` or `out_bp` by looking up their value in `table_in` or `table_out`. It can be implemented using `_mm256_permutevar_ps` in AVX2 or `vqtbl1q_u8` in NEON.

#### MulHash

A drawback of ntHash is that the efficient SIMD table lookup only works because it uses an alphabet of size 4. This means it does not work for general ASCII input. We introduce an alternative that we call *mulHash*, which replaces the table lookup of ntHash by *f* (*x*) = *C* · *x*, where *C* is a fixed random constant and the multiplication is wrapping over 32-bit integers. Although slightly slower, this can be easily computed for any input value *x*.

### 3.3 Sliding Window Minimum

The second step of our algorithm computes the position of the minimum *k*-mer hash in each sliding window of *w* hashes. A number of different approaches can be used for this, and pseudocode for each of the methods discussed can be found in Algorithm 3.

#### Naive

The simplest approach is to simply loop over the *w* values in each window independently. This takes *O*(*wn*) time, but can still be quite efficient when *w* is small by using vectorized instructions.

#### Monotone queue

An approach with better complexity is to use a *monotone queue*, which stores a non-decreasing subsequence of the *w* hashes, alongside their positions. Every time the window slides one to the right and we are about to push a new *k*-mer hash onto the right of the queue, we first remove any values larger than it, as they are “shadowed” by the new hash and can never be minimal anymore. The minimum of the window is then always the leftmost queue element. This data structure guarantees an amortized constant time update, but has many unpredictable branches due to removing between 0 and *w* values, which makes it costly in practice.

#### Rescan

Another approach used in bioinformatics is to only keep track of the minimum value and rescan the entire window of *w* values when the current minimum goes out of scope [17, 18]. While this algorithm does not guarantee a worst-case constant time update, it only branches when the minimum goes out of scope and hence is more predictable. This makes it more efficient in practice, especially since minimizers typically have a density of *O*(1*/w*) so that the *O*(*w*) rescan step takes amortized constant time per element.

To the best of our knowledge, most existing methods in bioinformatics use either a monotone queue or a rescan approach [11, 17, 18].

#### Two-stacks

Since our goal here is to compute *L* minima at the same time using vectorized instructions, we want to avoid any kind of data-dependent branches to ensure that the code path is the same for each chunk. A method for *online* sliding minima where elements may be added and removed at varying rates is the *two-stacks* method [10, 30], that is well-known in the competitive programming community^1^. Here, we only discuss a version where the number of elements remains constant at *w*.

##### Algorithm 3

Scalar implementations of the naive, queue and rescan sliding minimum methods, that return the position of the leftmost minimum value in each window of size *w*.

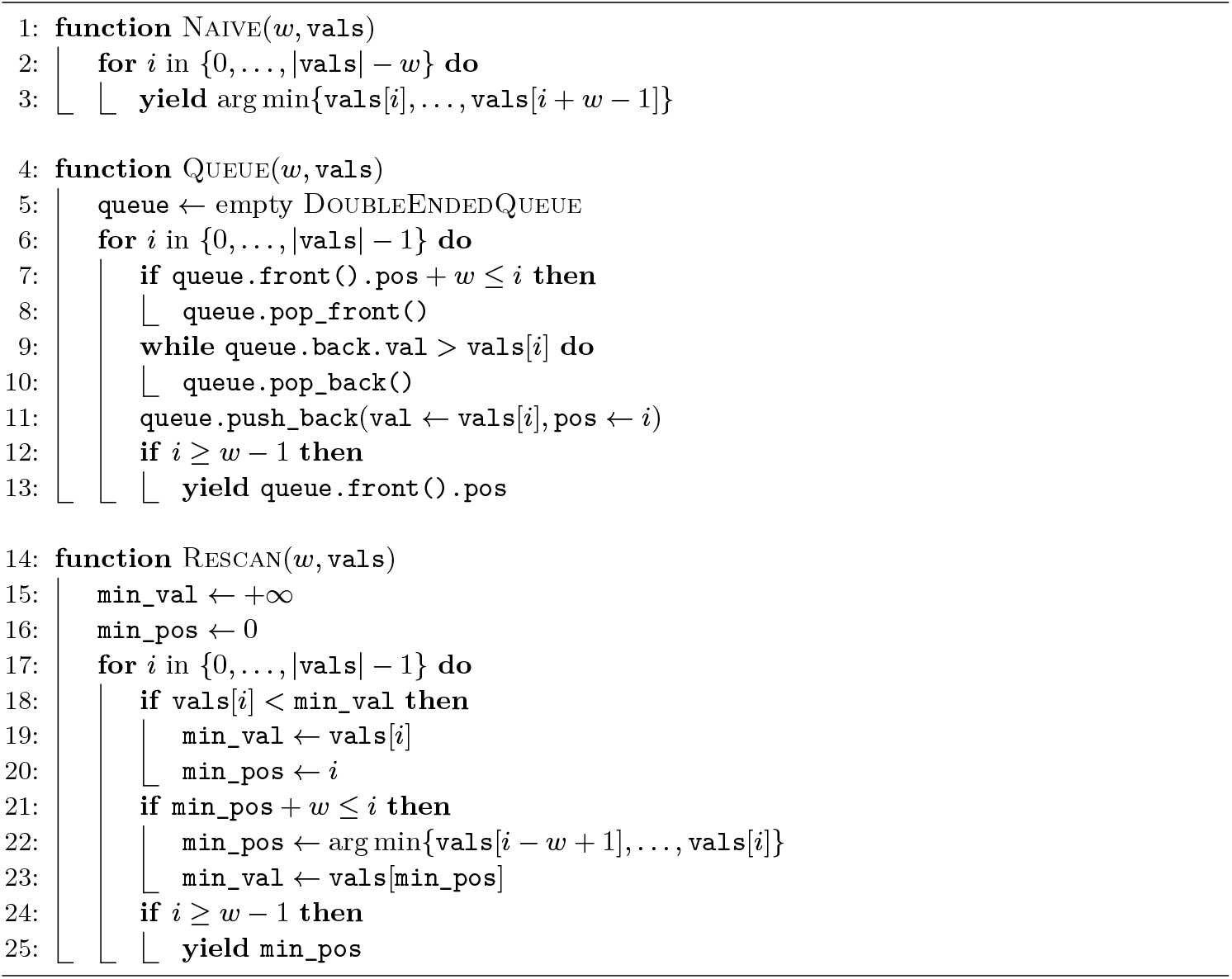

Conceptually, we first split the sequence of input *k*-mer hashes into blocks of size *w*, as shown in Figure 2. Then, we can compute both prefix minima and suffix minima of each block in *O*(*w*) per block, or *O*(1) amortized per input hash. Now, any window of size *w* can be split into a suffix of the previous block and a prefix of the current block (as highlighted in Figure 2), and we can return the minimum of the two corresponding suffix/prefix minima. When the window exactly coincides with a block (as shown on the right), the suffix and prefix minimum are equal.

**Figure 2.**
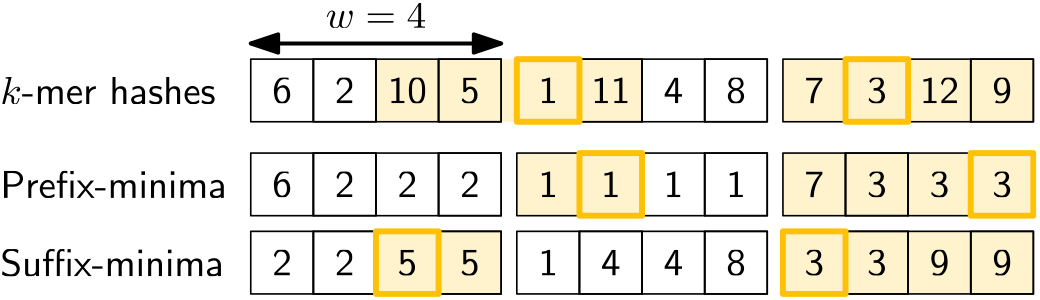
An example showing how sliding-window minima can be computed for windows of size *w* = 4. The input *k*-mer hashes are split into blocks of size *w*, and for each block, prefix and suffix minima are computed. Then, each window that overlaps two blocks can be split into a suffix and prefix, and the two corresponding minima (highlighted, 5 and 1) can be looked up. The minimum of these two is the minimum of the window. Windows that coincide with a block (as shown on the right) simply use the entire block itself as both prefix and suffix. In practice, we track the position of each minimum alongside its value.

In the implementation, Algorithm 4, the prefix minima are simply computed incrementally, while the suffix-minima are computed in batches after every block of *w* hashes has been filled. This way, only a single buffer of size *w* is needed, and the two cases above are unified.

The only branch in the algorithm triggers exactly every *w* iterations, and is thus completely data-independent. Thus, branches are both highly predictable, and the same across all *L* lanes.

##### Algorithm 4

Sliding minimum computation using a method based on the *two-stacks* algorithm. The input is an iterator over 32-bit hash values of the *k*-mers in each of the chunks as returned by ntHash (Algorithm 2), and the output is an iterator over the position of the leftmost minimum hash in each window of *w* hashes. Each operation can be vectorized to work on *L* lanes in parallel, each identified by a `LANE_IDX` ∈ [0, *L* − 1].

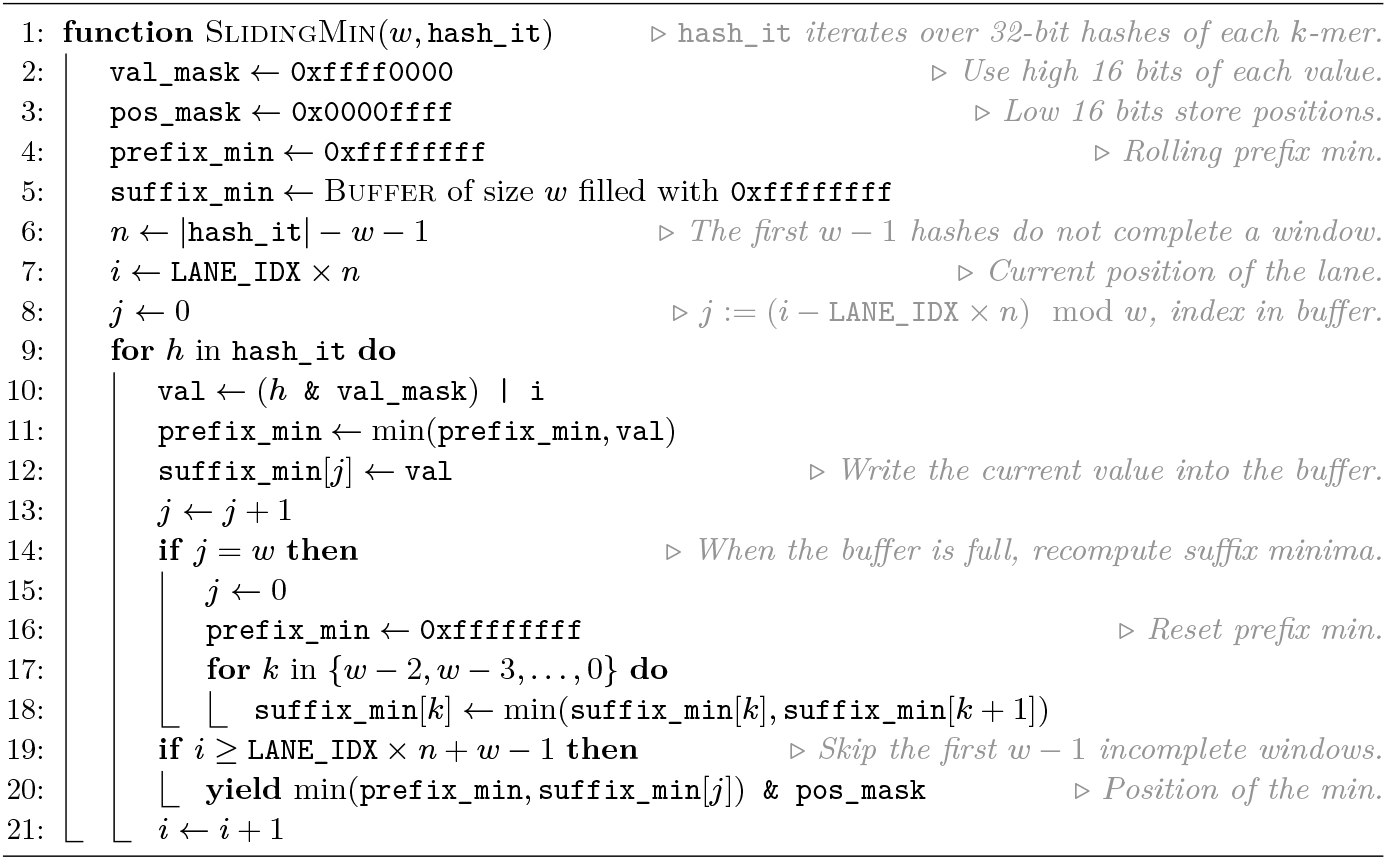

#### 16-bit hashes

To easily return the leftmost position of the minimum, instead of the minimum itself, we only use the upper 16 bits of each hash value, and store the position in the lower 16 bits. After taking the minimum, we mask out the high bits to obtain its position. Strings longer than 2^16^ characters can be processed in chunks of 2^16^. We note here that while using a 16-bit hash is usually not sufficient for *k*-mer indexing purposes, this *is* sufficiently good for selecting the minimum of a window when *w* ≪ 2^16^, which is indeed the case in bioinformatics applications, where usually *w* ≤ 100. Our library intentionally does not expose this hash to the user. Instead, a second, larger, hash can be computed afterwards for indexing purposes if needed.

### 3.4 Canonical Minimizers

One problem that arises when using minimizers in practice is that the DNA *strand* is often unknown. Thus, we do not know whether we are reading the *forward* (sense) or *reverse-complement* (antisense) strand, where the sequence is reversed and complementary bases `A ↔T` and `C↔ G` are used. When given a sequence, we would like to select the same minimizers in a strand-agnostic manner.

#### Canonical strand

Following [25], we define the *canonical* strand for each window of length *𝓁* as the strand where the count of `GT` bases (encoded values 3 and 2) is the highest, as shown in Algorithm 5. (In fact, any pair of bases can be chosen, as long as they are not complementary.) This count is in [0, *𝓁*] and when *𝓁* is odd there can be no tie between the two strands. In code, we instead compute the more symmetric `#GT − #AC = 2#GT` − *𝓁*, which is in [−*𝓁, 𝓁*], so that count *>* 0 defines the canonical strand. Then, we select the leftmost forward minimizer when the input window is canonical, and the rightmost reverse-complement minimizer when the input window is not canonical.

##### Algorithm 5

Counting `#GT − #AC`. When *𝓁* is odd, this can never be 0, and a window is canonical when the value is positive. The input is a stream over bases entering and leaving each *window*, where one is delayed by *𝓁* − 1 = *w* + *k* − 2 steps. Each operation can be vectorized to work on *L* lanes in parallel.

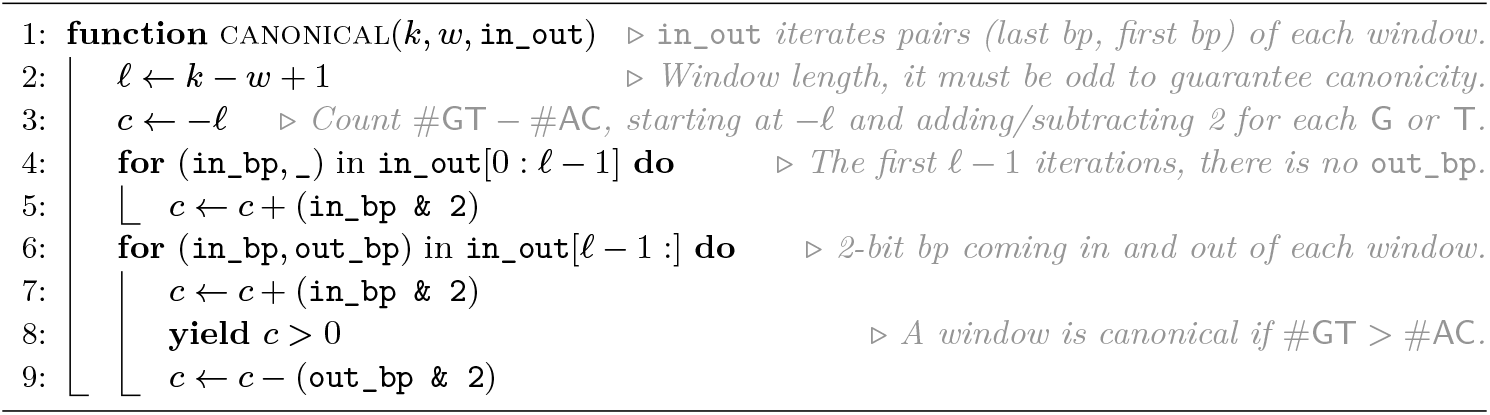

The benefit of this method over, say, determining the strand via the middle character (assuming again that *𝓁* is odd), is that the `GT` count is more stable across consecutive windows, since it varies by ±1. This way, the strandedness and thus the chosen minimizer is less likely to flip between adjacent windows.

#### Canonical ntHash

NtHash can be easily modified to compute both the forward and reverse-complement hash of each *k*-mer at the same time. Then, we can duplicate the sliding minimum algorithm to find the minimum of (*h*_fwd_(*kmer*[*i*]), *i*) and the *maximum* of (− *h*_rc_(*kmer*[*i*]), *i*) over each window. This way, ties in the forward direction are broken towards small *i* and ties in the reverse-complement direction are broken towards large *i*. To avoid selecting a new minimizer whenever the strand changes, we use a strand-independent *canonical* version of ntHash for both strands, defined as *h*_c_ = *h*_fwd_ + *h*_rc_ [14], and we do not distinguish which strand a minimizer *k*-mer was chosen from.

While this process does not assume *k* to be odd, this may be a useful additional assumption for further processing of the minimizers, so that the canonical representation of each *k*-mer itself can be indexed. Alternatively, *k*-mers could all be processed in the direction of the strand of the window they minimize, but this requires additional bookkeeping to store this direction, and would require duplicating *k*-mers when they minimize both forward and reverse-complement strands.

#### Canonical minimizers are not forward

One small drawback of this scheme is that it is not *forward*. Suppose a long window has many occurrences of the smallest minimizer, and that shifting the window one position changes its canonical strand. Then, the position of the sampled minimizer could jump backwards: from sampling the rightmost minimizer to sampling the leftmost minimizer. In practice, this does not seem to be a major limitation, both because it is rare and because downstream methods usually work fine on non-forward schemes anyway.

## 4 A SIMD algorithm

### SIMD

So far, our algorithms for iterating the input bases (Algorithm 1), hashing *k*-mers (Algorithm 2), and computing sliding window minima (Algorithm 4) are completely scalar. Now, we would like to use SIMD instructions to speed them up. With 256-bit AVX2 instructions, for example, we can process *L* = 8 *lanes* of 32-bit values at a time. A first approach could be to use this to compute the hash of *L* consecutive minimizer values at the same time, and then to compute the minimizer of the *L* new windows all at once. Unfortunately, this is tricky due to the sequential nature of rolling hashing and sliding window minima.

### Chunks

Instead of processing *consecutive k*-mers in parallel, we choose to split the input sequence into *L* equally long *chunks* that we process in parallel. This way, we compute one hash of each chunk in parallel, and then compute one minimizer position of a window of each chunk in parallel as well. This works, because the computations on the chunks are completely independent of each other. We let adjacent chunks overlap by *𝓁* −1 characters, so that each window is fully contained in exactly one chunk. The total number of windows may not be divisible by *L*. In that case, we round up the chunk length and return the number of elements that was added as padding, so that these can be removed later. For simplicity, we omit that case from the code snippets.

### 4.1 Gathering the input

The parts that need special attention for SIMD instructions are the parallel reading of the input sequence, and the parallel deduplicating of the output minimizer positions (Section 4.2).

The first step is to read the actual bases. Since memory access instructions are relatively slow^2^ compared to the SIMD operations we do on them, we cannot afford to read from each chunk one base or one byte at a time. We could read 32 bits at a time from each chunk using SIMD `gather` instructions, but these are relatively slow. Instead^3^, we read a full SIMD register of 256 bits worth of data at a time from each chunk, as shown in Algorithm 6. Then, we *transpose*^4,5^ *this matrix so that we obtain L* SIMD registers, where the first contains the upcoming `u32` for each chunk, the second contains the next `u32` for each lane, and so on. Then, we use this transposed matrix as a buffer, and use it for the next 128 bases. After each 16 bases, we shift to the next `u32×8` from it.

#### Algorithm 6

Pseudocode for splitting the input sequence into *L* chunks and iterating each chunk in parallel. The input sequence is assumed to be bitpacked, and indexed by bytes. Yields an iterator over *L* lanes of `u32` values.

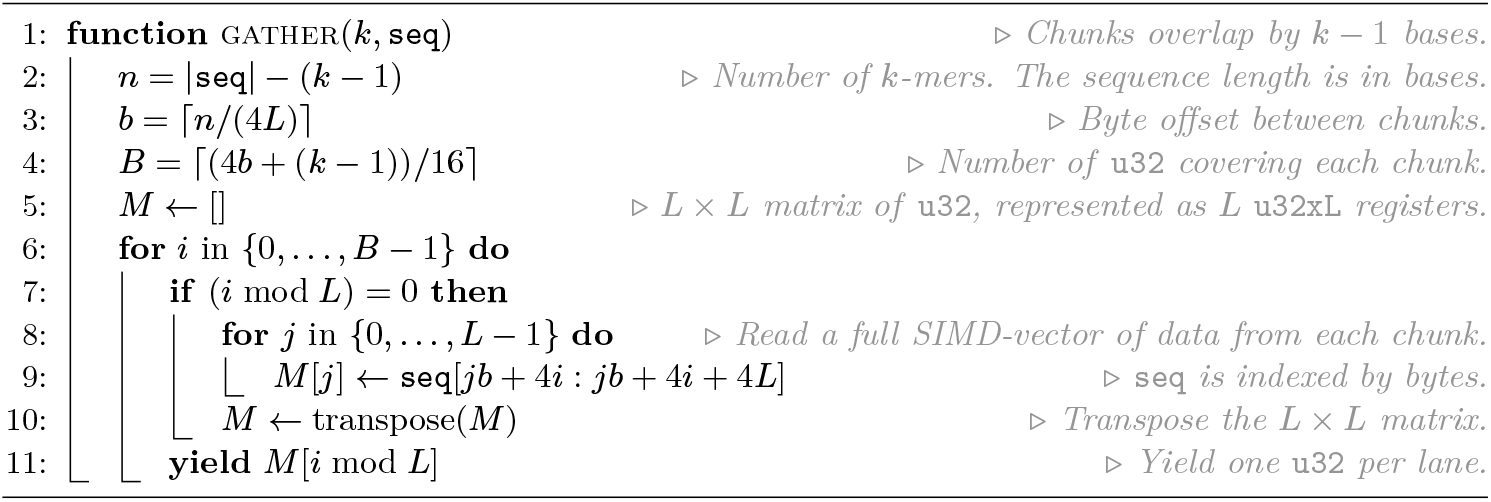

### 4.2 Collecting and Deduplicating Positions

#### Transpose

The last step of the minimizer algorithm is to deduplicate the results, since adjacent windows often share the same minimizer position. In practice, deduplicating works best when the data to be duplicated is linear in memory. But the output of the vectorized sliding-window-minimum gives a `u32×8` containing the position of one minimizer of each chunk. Thus, every *L* iterations we reuse the matrix transpose to obtain a `u32×8` for each chunk, containing *L* consecutive minimizer positions^6^.

#### Dedup

We deduplicate each lane using the technique of [16]. This compares each element to the previous one, and compares the first element to the last minimizer of the window before. The distinct elements are then *shuffled* to the front of the SIMD vector using a lookup table and appended to a buffer for each lane (Figure 4). We end by concatenating all the per-lane buffers into a single vector of minimizer positions, and make sure to avoid duplicates between the end and start of adjacent lanes.

**Figure 3.**
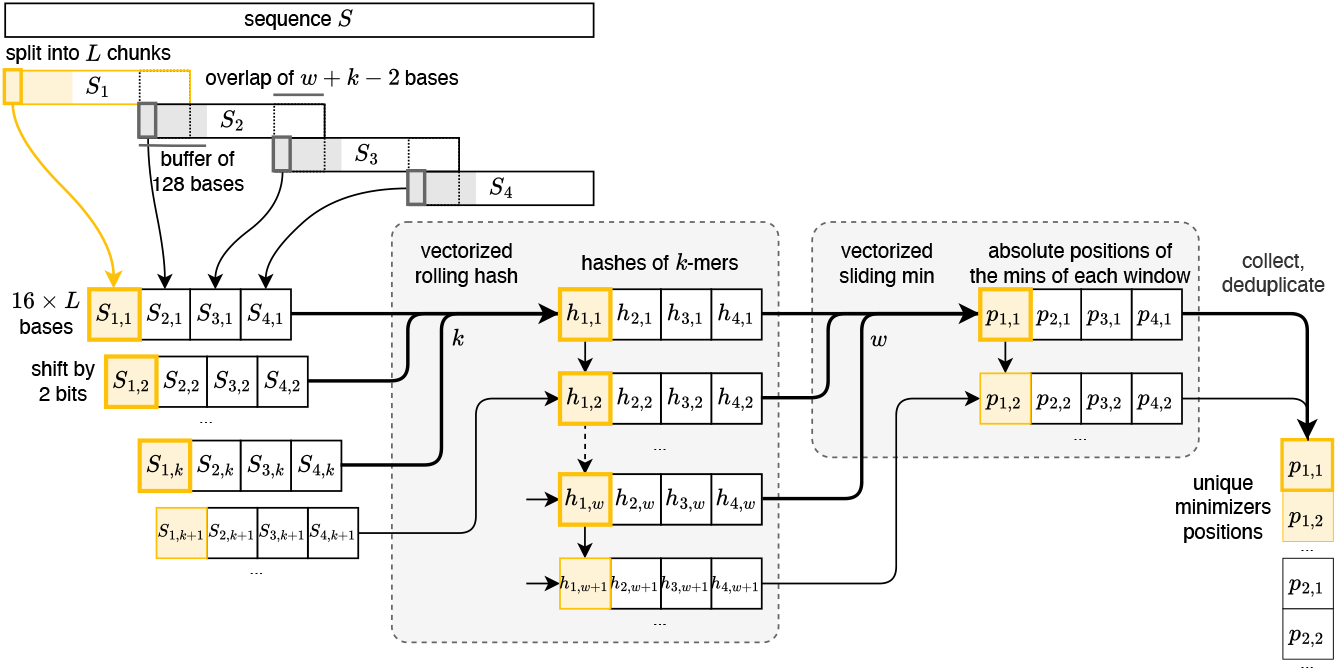
High-level view of the vectorized computation of minimizers using the building blocks presented in Section 3. The sequence is first split into *L* chunks (in this example *L* = 4) that are processed in parallel. As described in Section 4.1, we load 128 bases at a time for each chunk, from which we extract 16 bases *S*_*i,j*_ which are gathered in a SIMD register (one lane for each chunk, 32 bits per lane). This SIMD register is used to iterate over each lane by shifting and masking 2 bits at a time, and is passed as input to a vectorized rolling hash function that computes a hash *h*_*i,j*_ for the *k*-mers in each lane. The absolute position *p*_*i,j*_ of the minimum is then computed over a sliding window of *w* vectors of hashes for each lane. The positions are finally reordered to match the order of the original sequence and deduplicated to keep a unique occurrence of each position. At every step of the computation, the lane corresponding to the first chunk is highlighted in yellow. A bold outline indicates the bases in the first *k*-mer and the hashes corresponding to the first window.

**Figure 4.**
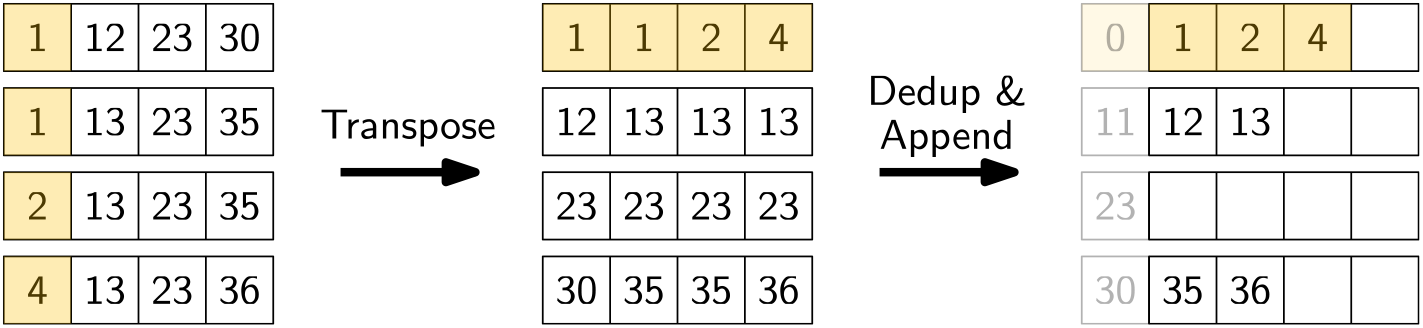
The SIMD version of the sliding window minimum algorithm produces minimizer positions for *L* chunks at a time. For example, the top row on the left indicates a SIMD vector containing the minimizer position of a window in each of the *L* = 4 chunks. In the end, we want to return a single “flat” vector. Thus, we have to “deinterleave” the *L* lanes. First, we collect *L* minimizer positions of every chunk (the matrix on the left). Then, we transpose this matrix, so that the resulting SIMD vectors each correspond to a single chunk, with values corresponding to the first chunk highlighted. These are then compared with their preceding element (including the previous minimizer position, as shown in grey), and distinct elements are shuffled to the front. These positions are accumulated in a separate buffer for each chunk, and these buffers are finally concatenated into a single flat vector.

#### Super-*k*-mers

The deduplication can be amended to also find super-*k*-mers, which are sequences of consecutive windows sharing the same minimizer position. After comparing adjacent minimizer positions, we obtain a mask that determines the shuffle instruction to apply. Normally we shuffle the 32-bit minimizer positions directly. Instead, we can mask out the upper 16 bits and store there the index of its window. If we then shuffle those values, we obtain for each minimizer its position in the input text, and the position of the first window where this *k*-mer became a minimizer. This information is sufficient to recover all super-*k*-mers, and sequences longer than 2^16^ bp can be either split-up, or else it is easy to detect manually when the values wrapped.

## 5 Experimental Evaluation

Our code is available in the `simd-minimizers` crate that can be found at https://github.com/rust-seq/simd-minimizers. Part of the code was extracted into a separate library, packed-seq, for easy reuse in other projects, which is available at https://github.com/rust-seq/packed-seq. Both libraries support both AVX2 and NEON instruction sets, and we will now look at their performance. The code to reproduce the experiments is available in the `simd-minimizers`-bench subdirectory.

The experiments were run on an Intel Core i7-10750H with 6 cores with AVX2 running at a fixed frequency of 2.6GHz with hyperthreading disabled and cache sizes of 32 KiB (L1), 256 KiB (L2), and 12 MiB shared L3. Code is compiled with `rustc 1.88.0-nightly` ^7^. As input, we use a fixed random string of 10^8^ bases, depending on the method encoded either as ASCII or as packed representation. Reported timings are the median of five runs and shown in nanoseconds per base.

In our experiments, we use parameter values for *w* and *k* as used by Kraken2 [31] (5, 31), SSHash [27] (11, 21), and Minimap2 [17] (19, 19).

Tables 3 and 4 in Section A show equivalent results for the NEON architecture.

### Incremental time usage

Table 1 shows the time usage for various incremental subsets of our method. To start, iterating the 8 chunks of the input and summing all bases takes 0.15 ns/bp. Appending all `u32×8` SIMD vectors containing the bases to a vector takes 0.30 ns/bp, indicating that writing to memory induces some overhead. Collecting the second *k* − 1-delayed stream of characters that leave the *k*-mer (by adding them to the non-delayed stream) has no additional overhead. Computing ntHash only takes 0.02ns/bp extra. The sliding window computation nearly triples the total time. Collecting the minimizer positions to a linear vector (i.e., transposing matrices and writing output for each of the 8 chunks) again incurs 50% overhead, ad deduplicating them actually again adds some time.

**Table 1.** Total time per base taken when incrementally including more steps of the implementation, for (*w, k*) = (11, 21).

| Part | ns/bp |
| --- | --- |
| Iterate the bases | 0.15 |
| + collect to vector | 0.30 |
| + iterate the delayed bases | 0.30 |
| + ntHash | 0.32 |
| + sliding window min | 0.90 |
| + collect | 1.48 |
| + dedup | **1.61** |
| + canonical nthash | 1.04 |
| + canonical strand | 1.53 |
| + collect | 2.03 |
| + dedup | **2.20** |

Going back a step, using canonical ntHash instead of forward ntHash takes 0.14 ns/bp extra, and determining the canonical strand (via a third *𝓁*-delayed stream and counting `GT` bases) takes another 0.49 ns/bp. As before, collecting and deduplicating are slow and add around 0.70ns/bp.

In conclusion, we see that iterating the chunks of the input and determining minimizers is quite fast, but that a lot of time must then be spent to “deinterleave” the output into a linear stream. As can be expected, canonical minimizers are slower to compute than forward minimizers, but the overhead is less than 50%, which seems quite low given that the ntHash and sliding window minimum computation are duplicated and a canonical-strand computation is added.

### Full comparison

We compare against the `minimizer-iter` crate (v1.2.1) [20], which implements a queue-based sliding window minimum using wyhash [32] and also supports canonical minimizers. For an additional comparison, we optimized an implementation of the remap method with ntHash based on a code snippet by Daniel Liu [18].

Results are in Table 2 and Figure 5. `minimizer-iter` takes around 26ns/bp for forward and 33ns/bp for canonical minimizers, and its runtime does not depend much on *w* and *k*, because the popping from the queue is unpredictable regardless of *w*. Rescan starts out at 11.7 ns/bp for *w* = 5 and gets significantly faster as *w* increases, converging to around 5 ns/bp for *w* ≫ 100. This is explained by the fact that rescan has a branch miss every time the current minimizer falls out of the window, which happens for roughly half the minimizers at a rate of 1*/*(*w* + 1). Thus, as *w* increases, the method becomes more predictable and branch misses go down. Our method, `simd-minimizers`, runs around 1.61 ns/bp for forward and 2.20 ns/bp for canonical minimizers when given packed input, and therefore is 3.4× to 6.8 × faster than the rescan method.

**Table 2.**
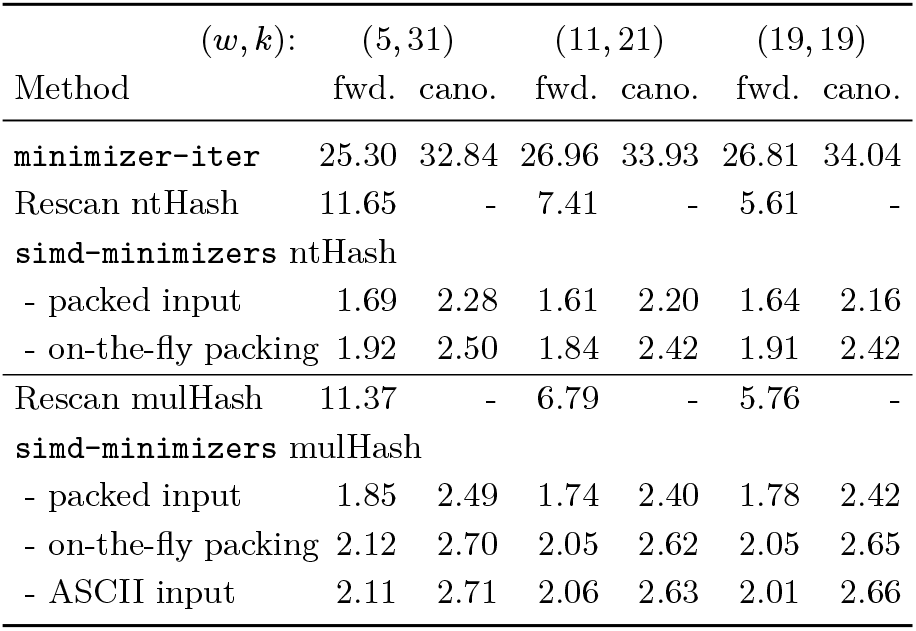
Comparison of our `simd-minimizers` implementation against `minimizer-iter` [20] and a rescan implementation based on [18]. Times in ns/bp are shown for both forward and canonical minimizers (where supported), and for various (*w, k*) tuples. For our library, we test both ntHash and mulHash with multiple encodings of the input DNA: 1) 2-bit packed, 2) ASCII-`ACGT` that is packed on-the-fly, and 3) plain ASCII (for mulHash only).

**Figure 5.**
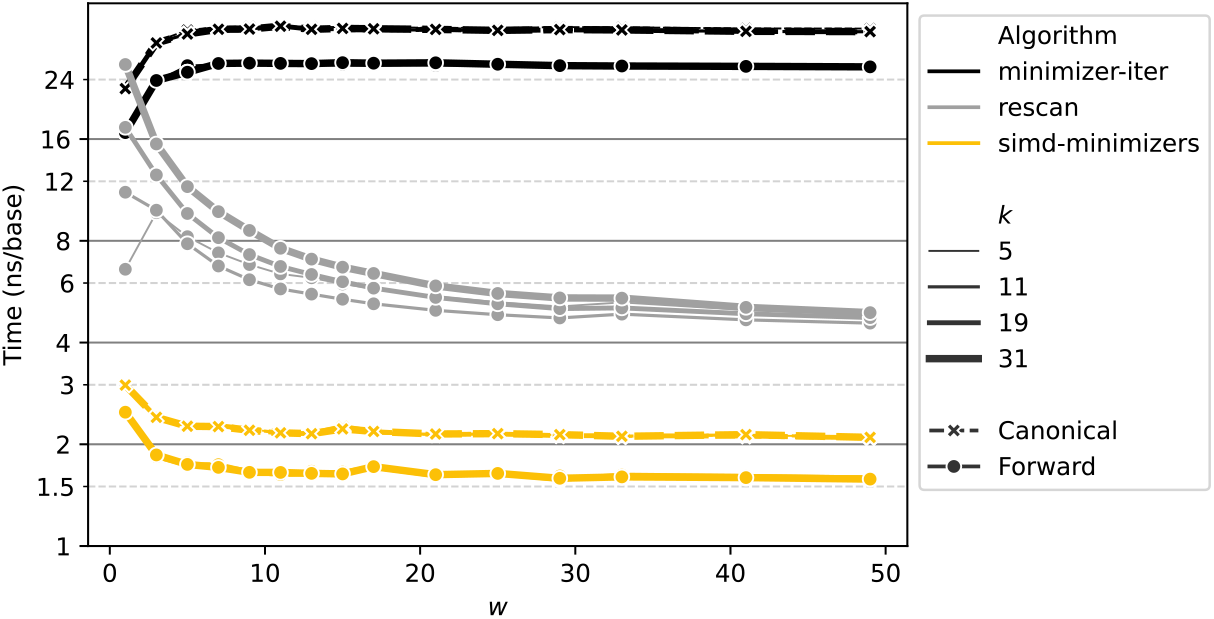
Running time (logarithmic) of `minimizer-iter`, rescan, and `simd-minimizers` for different values of *w* and *k*. For `minimizer-iter` and `simd-minimizers`, runtime is nearly independent of *k*, while both are slower when computing canonical minimizers (indicated with crosses).

**Figure 6.**
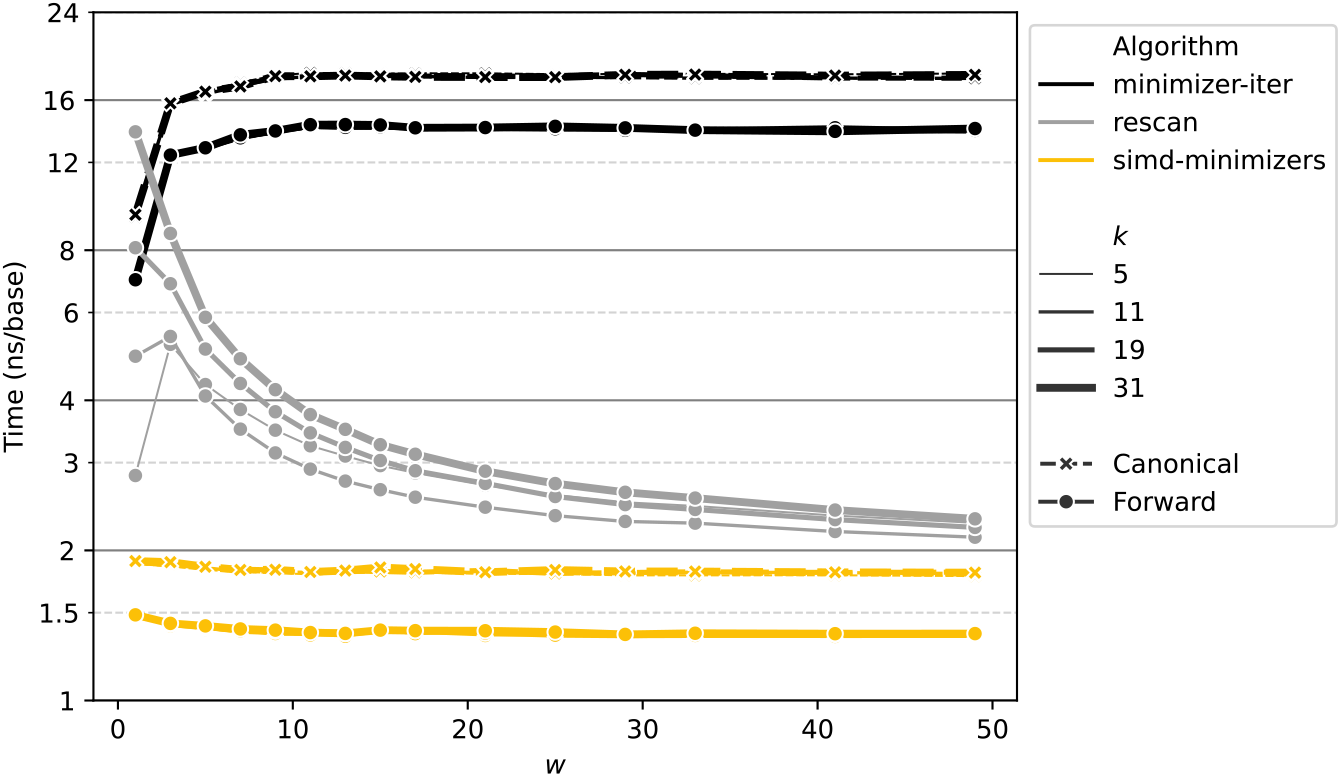
Running time of `minimizer-iter`, rescan and `simd-minimizers` on NEON architecture for different values of *w* and *k*.

In Figure 5, we see that `simd-minimizers`’ performance is mostly independent of *k* and *w* since it is mostly data-independent. Only for small *w* ≤ 5 it is slightly slower due to the larger number of minimizers and hence larger size of the output.

As we use SIMD with 8 lanes, we could in theory expect up to 8× speedup. In practice this is hard to reach because of constant overhead and because the overhead to work well with SIMD in the first place. In particular for large *w*, rescan benefits from very predictable and simple code and only outputs unique minimizer positions, making it very efficient. In SIMD, on the other hand, we use a data-independent algorithm, and output the minimizer position for every single window, which then has to be deduplicated. Thus, it is nice to see that even for large *w*, our method is over 3× faster, despite this overhead.

### ASCII input and mulHash

Apart from taking bit-packed input, `simd-minimizers` also works on ASCII-encoded DNA sequences of `ACTG` characters directly, which are then packed into values {0, 1, 2, 3} for ntHash (in that order) during iteration. This is around 0.25ns/bp slower, mostly because of the larger size of the unpacked input.

The mulHash variant is around 0.20ns/bp slower again, but works for any ASCII input. Performance on 100 MB of the Pizza&Chili corpus [5] English^8^ and Sources^9^ datasets is nearly identical to performance on the random DNA shown in Table 2.

### Human genome

We also run `simd-minimizers` on the chromosomes of a human genome (T2T-CHM13v2.0^10^ [24]), of total size 3.2 Gbp. Here, computing forward minimizers takes 5.19 seconds, and canonical minimizers takes 6.71 seconds for (*w, k*) = (11, 21), which corresponds 1.67 and 2.15 ns/bp, which is within a few percent of Table 2.

### Density

For (*w, k*) = (11, 21), we get a density of 0.173 for forward minimizers, and 0.167 for canonical minimizers, which are both close to the expected density of 2*/*(*w* +1) = 1*/*6 ≈ 0.167. Changing to (*w, k*) = (19, 19), we get density 0.098 for forward minimizers and 0.100 for canonical minimizers, both close to the expected density of 2*/*(19 + 1) = 1*/*10 = 0.1. Thus, we see that in practice, ntHash is a sufficiently random hash function to approach the expected density of random minimizers with a perfectly uniform random hash function.

### Multithreading

We test throughput in a multithreaded setting, by using 6 threads to process the 25 chromosomes (22, X, Y, and mitochondrion) in parallel. This way, processing takes 0.97 s (forward) and 1.27 s (canonical) when the input data is already loaded into memory, showing slightly above 5× speedup. This is just below 6× speedup as the chromosomes don’t perfectly partition the data into 6× equal parts, so that some threads finish before others.

For applications, we recommend to simply call our library in parallel from multiple threads as needed.

## 6 Conclusions and Future Work

Our library `simd-minimizers` computes minimizer positions 3.4× to 6.8× faster than other methods. Using the library, only a single function call is needed to obtain the list of (canonical) minimizer positions, taking as input the (packed) DNA sequence and the parameters *k* and *w*. General ASCII input are also supported, allowing use cases such as sketching protein sequences. We hope that the community will adopt `simd-minimizers` as the standard library to compute random minimizers.

### Future work

We chose to use the data-independent two-stacks method as the core of our algorithm, that returns the minimizer position of every window and then requires deduplication. Given the promising performance of rescan for large *w* in Table 2, an interesting alternative could be to go the opposite way and speed up the rescan step. This is particularly relevant when minimizers are sparse, as the number of branches may be small enough for linear scans to benefit from SIMD. Accelerating linear scans could also be useful for smaller inputs such as short reads, where splitting into 8 chunks may not be very efficient.

Another approach would be to use 512-bit AVX512 instructions and process 16 lanes in parallel. In theory that could be another 2× faster, but in practice the collecting and deduplicating of values may become an even larger bottleneck. In simple experiments, without further profiling and optimizing the code, it is in fact 20% *slower*.

Additionally, implementing low-density schemes like the open-closed mod-minimizer [7, 9] would be a valuable extension.

## Supplementary Material

*Software*: https://github.com/rust-seq/simd-minimizers

*Software*: https://github.com/rust-seq/packed-seq

## Acknowledgements

We thank the anonymous reviewers for their comments that lead to a significantly improved presentation, as well as for their suggestion to apply a matrix transpose to the input, which improved performances.

## A Results for NEON Architecture

The NEON experiments are run on a performance core on an Apple M1 chip with 4 efficiency and 4 performance cores. The performance core runs at 3.2GHz and has 128KiB of L1 cache, 12MiB of shared L2 cache (for the performance cores) and 8MiB of shared L3 cache (for the whole system).

**Table 3.**
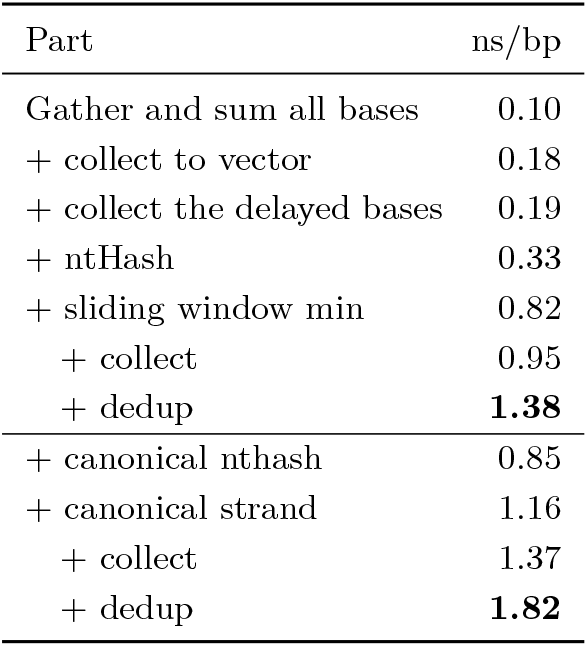
Time per base taken when adding steps of the implementation, for (*w, k*) = (11, 21) on NEON architecture.

**Table 4.** Comparison of our `simd-minimizers` implementation against `minimizer-iter` [20] and a rescan implementation based on [18]. Times in ns/bp are shown for both forward and canonical minimizers (where supported), and for various (*w, k*) tuples, on NEON architecture.

| | $(w, k): (5, 31)$ | | $(11, 21)$ | | $(19, 19)$ | |
| --- | --- | --- | --- | --- | --- | --- |
| Method | fwd. | cano. | fwd. | cano. | fwd. | cano. |
| `minimizer-iter` | 12.89 | 16.60 | 14.26 | 18.08 | 14.20 | 18.04 |
| Rescan ntHash | 5.95 | - | 3.92 | - | 2.82 | - |
| `simd-minimizers` ntHash |  |  |  |  |  |  |
| - packed input | 1.42 | 1.85 | 1.38 | 1.82 | 1.38 | 1.83 |
| - on-the-fly packing | 1.66 | 2.07 | 1.58 | 1.99 | 1.59 | 2.02 |
| Rescan mulHash | 6.94 | - | 3.93 | - | 3.42 | - |
| `simd-minimizers` mulHash |  |  |  |  |  |  |
| - packed input | 1.97 | 4.34 | 1.91 | 4.27 | 1.93 | 4.26 |
| - on-the-fly packing | 2.11 | 4.37 | 2.04 | 4.41 | 2.03 | 4.30 |
| - ASCII input | 2.12 | 4.39 | 2.06 | 4.44 | 2.04 | 4.29 |

## Footnotes

1 https://codeforces.com/blog/entry/71687

2 https://uops.info/html-instr/VPGATHERDD_YMM_VSIB_YMM_YMM.html

3 As suggested by one of the anonymous reviewers, for which we thank them.

4 https://stackoverflow.com/questions/25622745/transpose-an-8x8-float-using-avx-avx2

5 https://github.com/rust-seq/simd-minimizers/blob/master/simd-minimizers/src/intrinsics/transpose.rs

6 https://github.com/rust-seq/simd-minimizers/blob/master/simd-minimizers/src/collect.rs

7 We have made sure that all layers of iterators are inlined into single function. This is usually needed for optimal performance. Even then, small changes to the generated code (by changing the source, or compiler version) can impact performance by as much as 20%.

8 https://pizzachili.dcc.uchile.cl/texts/nlang/

9 https://pizzachili.dcc.uchile.cl/texts/code/

10 Available at https://github.com/marbl/CHM13

## References

1 Gaëtan Benoit, Sébastien Raguideau, Robert James, Adam M Phillippy, Rayan Chikhi, and Christopher Quince. High-quality metagenome assembly from long accurate reads with metamdbg. Nature Biotechnology, pages 1–6, 2024. doi:10.1038/s41587-023-01983-6.

2 Jonathan D. Cohen. Recursive hashing functions for n-grams. ACM Trans. Inf. Syst., 15(3):291–320, July 1997. doi:10.1145/256163.256168.

3 Sebastian Deorowicz, Marek Kokot, Szymon Grabowski, and Agnieszka Debudaj-Grabysz. KMC 2: fast and resource-frugal k-mer counting. Bioinformatics, 31(10):1569–1576, January 2015. doi:10.1093/bioinformatics/btv022.

4 Baris Ekim, Bonnie Berger, and Rayan Chikhi. Minimizer-space de bruijn graphs: Wholegenome assembly of long reads in minutes on a personal computer. Cell Systems, 12(10):958–968.e6, 2021. URL: https://www.sciencedirect.com/science/article/pii/S240547122100332X, doi:10.1016/j.cels.2021.08.009.

5 Paolo Ferragina and Gonzalo Navarro. The Pizza&Chili Corpus. https://pizzachili.dcc.uchile.cl/texts.html, 2007.

6 Shay Golan, Ido Tziony, Matan Kraus, Yaron Orenstein, and Arseny Shur. Generating low-density minimizers. bioRxiv, November 2024. doi:10.1101/2024.10.28.620726.

7 Ragnar Groot Koerkamp, Daniel Liu, and Giulio Ermanno Pibiri. The open-closed mod-minimizer algorithm. bioRxiv, 2024. doi:10.1101/2024.11.02.621600.

8 Ragnar Groot Koerkamp and Igor Martayan. SimdMinimizers: Computing random minimizers, fast. bioRxiv, 2025. doi:10.1101/2024.11.02.621600.

9 Ragnar Groot Koerkamp and Giulio Ermanno Pibiri. The mod-minimizer: A Simple and Efficient Sampling Algorithm for Long k-mers. In Solon P. Pissis and Wing-Kin Sung, editors, 24th International Workshop on Algorithms in Bioinformatics (WABI 2024), volume 312 of Leibniz International Proceedings in Informatics (LIPIcs), pages 11:1–11:23, Dagstuhl, Germany, 2024. Schloss Dagstuhl – Leibniz-Zentrum für Informatik. URL: https://drops.dagstuhl.de/entities/document/10.4230/LIPIcs.WABI.2024.11, doi:10.4230/LIPIcs.WABI.2024.11.

10 Martin Hirzel, Scott Schneider, and Kanat Tangwongsan. Sliding-window aggregation algorithms: Tutorial. In Proceedings of the 11th ACM International Conference on Distributed and Event-based Systems, DEBS 2017, Barcelona, Spain, June 19-23, 2017, pages 11–14. ACM, 2017. doi:10.1145/3093742.3095107.

11 Guillaume Holley and Páll Melsted. Bifrost: highly parallel construction and indexing of colored and compacted de bruijn graphs. Genome biology, 21:1–20, 2020.

12 Chirag Jain, Arang Rhie, Nancy F Hansen, Sergey Koren, and Adam M Phillippy. Long-read mapping to repetitive reference sequences using winnowmap2. Nature Methods, 19(6):705–710, 2022. doi:10.1038/s41592-022-01457-8.

13 Richard M. Karp and Michael O. Rabin. Efficient randomized pattern-matching algorithms. IBM Journal of Research and Development, 31(2):249–260, 1987. doi:10.1147/rd.312.0249.

14 Parham Kazemi, Johnathan Wong, Vladimir Nikolić, Hamid Mohamadi, René L Warren, and Inanç Birol. nthash2: recursive spaced seed hashing for nucleotide sequences. Bioinformatics, 38(20):4812–4813, August 2022. doi:10.1093/bioinformatics/btac564.

15 Bryce Kille, Ragnar Groot Koerkamp, Drake McAdams, Alan Liu, and Todd J Treangen. A near-tight lower bound on the density of forward sampling schemes. Bioinformatics, 41(1):btae736, December 2024. doi:10.1093/bioinformatics/btae736.

16 Daniel Lemire. Removing duplicates from lists quickly. https://lemire.me/blog/2017/04/10/removing-duplicates-from-lists-quickly/, April 2017.

17 Heng Li. Minimap2: pairwise alignment for nucleotide sequences. Bioinformatics, 34(18):3094– 3100, May 2018. doi:10.1093/bioinformatics/bty191.

18 Daniel Liu. minimizer.rs re-scan implementation. https://gist.github.com/Daniel-Liu-c0deb0t/7078ebca04569068f15507aa856be6e8, July 2023.

19 Camille Marchet, Mael Kerbiriou, and Antoine Limasset. Blight: efficient exact associative structure for k-mers. Bioinformatics, 37(18):2858–2865, April 2021. doi:10.1093/bioinformatics/btab217.

20 Igor Martayan. minimizer-iter: Iterate over minimizers of a DNA sequence. https://github.com/rust-seq/minimizer-iter.

21 Igor Martayan, Lucas Robidou, Yoshihiro Shibuya, and Antoine Limasset. Hyper-k-mers: efficient streaming k-mers representation. bioRxiv, November 2024. doi:10.1101/2024.11.06.620789.

22 Guillaume Marçais, C S Elder, and Carl Kingsford. k-nonical space: sketching with reverse complements. Bioinformatics, 40(11):btae629, October 2024. doi:10.1093/bioinformatics/btae629.

23 Hamid Mohamadi, Justin Chu, Benjamin P. Vandervalk, and Inanc Birol. nthash: recursive nucleotide hashing. Bioinformatics, 32(22):3492–3494, July 2016. doi:10.1093/bioinformatics/btw397.

24 Sergey Nurk, Sergey Koren, Arang Rhie, Mikko Rautiainen, Andrey V. Bzikadze, Alla Mikheenko, Mitchell R. Vollger, Nicolas Altemose, Lev Uralsky, Ariel Gershman, Sergey Aganezov, Savannah J. Hoyt, Mark Diekhans, Glennis A. Logsdon, Michael Alonge, Stylianos E. Antonarakis, Matthew Borchers, Gerard G. Bouffard, Shelise Y. Brooks, Gina V. Caldas, Nae-Chyun Chen, Haoyu Cheng, Chen-Shan Chin, William Chow, Leonardo G. de Lima, Philip C. Dishuck, Richard Durbin, Tatiana Dvorkina, Ian T. Fiddes, Giulio Formenti, Robert S. Fulton, Arkarachai Fungtammasan, Erik Garrison, Patrick G. S. Grady, Tina A. Graves-Lindsay, Ira M. Hall, Nancy F. Hansen, Gabrielle A. Hartley, Marina Haukness, Kerstin Howe, Michael W. Hunkapiller, Chirag Jain, Miten Jain, Erich D. Jarvis, Peter Kerpedjiev, Melanie Kirsche, Mikhail Kolmogorov, Jonas Korlach, Milinn Kremitzki, Heng Li, Valerie V. Maduro, Tobias Marschall, Ann M. McCartney, Jennifer McDaniel, Danny E. Miller, James C. Mullikin, Eugene W. Myers, Nathan D. Olson, Benedict Paten, Paul Peluso, Pavel A. Pevzner, David Porubsky, Tamara Potapova, Evgeny I. Rogaev, Jeffrey A. Rosenfeld, Steven L. Salzberg, Valerie A. Schneider, Fritz J. Sedlazeck, Kishwar Shafin, Colin J. Shew, Alaina Shumate, Ying Sims, Arian F. A. Smit, Daniela C. Soto, Ivan Sović, Jessica M. Storer, Aaron Streets, Beth A. Sullivan, Françoise Thibaud-Nissen, James Torrance, Justin Wagner, Brian P. Walenz, Aaron Wenger, Jonathan M. D. Wood, Chunlin Xiao, Stephanie M. Yan, Alice C. Young, Samantha Zarate, Urvashi Surti, Rajiv C. McCoy, Megan Y. Dennis, Ivan A. Alexandrov, Jennifer L. Gerton, Rachel J. O’Neill, Winston Timp, Justin M. Zook, Michael C. Schatz, Evan E. Eichler, Karen H. Miga, and Adam M. Phillippy. The complete sequence of a human genome. Science, 376(6588):44–53, April 2022. URL: http://dx.doi.org/10.1126/science.abj6987, doi:10.1126/science.abj6987.

25 Chenxu Pan and Knut Reinert. A simple refined dna minimizer operator enables 2-fold faster computation. Bioinformatics, 40(2):btae045, January 2024. doi:10.1093/bioinformatics/btae045.

26 David Pellow, Lianrong Pu, Baris Ekim, Lior Kotlar, Bonnie Berger, Ron Shamir, and Yaron Orenstein. Efficient minimizer orders for large values of k using minimum decycling sets. Genome Research, 33(7):1154–1161, 2023. URL: https://www.genome.org/cgi/doi/10.1101/gr.277644.123, doi:10.1101/gr.277644.123.

27 Giulio Ermanno Pibiri. Sparse and skew hashing of k-mers. Bioinformatics, 38(Supplement_1):i185–i194, June 2022. doi:10.1093/bioinformatics/btac245.

28 Michael Roberts, Wayne Hayes, Brian R. Hunt, Stephen M. Mount, and James A. Yorke. Reducing storage requirements for biological sequence comparison. Bioinformatics, 20(18):3363– 3369, July 2004. doi:10.1093/bioinformatics/bth408.

29 Saul Schleimer, Daniel S. Wilkerson, and Alex Aiken. Winnowing: local algorithms for document fingerprinting. In Proceedings of the 2003 ACM SIGMOD international conference on Management of data, SIGMOD ‘03, pages 76–85, New York, NY, USA, June 2003. Association for Computing Machinery. URL: https://dl.acm.org/doi/10.1145/872757.872770, doi: 10.1145/872757.872770.

30 Georgios Theodorakis, Alexandros Koliousis, Peter R. Pietzuch, and Holger Pirk. Hammer slide: Work- and cpu-efficient streaming window aggregation. In Rajesh Bordawekar and Tirthankar Lahiri, editors, International Workshop on Accelerating Analytics and Data Management Systems Using Modern Processor and Storage Architectures, ADMS@VLDB 2018, Rio de Janeiro, Brazil, August 27, 2018, pages 34–41, 2018. URL: http://www.adms-conf.org/2018-camera-ready/SIMDWindowPaper_ADMS%2718.pdf.

31 Derrick E Wood, Jennifer Lu, and Ben Langmead. Improved metagenomic analysis with Kraken 2. Genome biology, 20:1–13, 2019. doi:10.1186/s13059-019-1891-0.

32 Wang Yi and Diego Barrios Romero. wyhash-rs, fast portable non-cryptographic hashing algorithm. https://github.com/eldruin/wyhash-rs.

33 Hongyu Zheng, Carl Kingsford, and Guillaume Marçais. Improved design and analysis of practical minimizers. Bioinformatics, 36(Supplement_1):i119–i127, July 2020. doi:10.1093/bioinformatics/btaa472.

